# Integrated ligand and structure-based investigation of structural requirements for silent information regulator [SIRT1] activation

**DOI:** 10.1101/481952

**Authors:** Amit K. Gupta, Sun Choi

**Affiliations:** National Leading Research Laboratory of Molecular Modeling & Drug Design, College of Pharmacy and Graduate School of Pharmaceutical Sciences, and Global Top 5 Research Program, Ewha Womans University, Seoul 03760, Republic of Korea.

**Keywords:** SIRT1 activators, imidazothiazole and oxazolopyridine derivatives, 3D QSAR, 2D QSAR, CoMFA and CoMSIA

## Abstract

A series of imidazothiazole and oxazolopyridine derivatives as human silent information regulator (SIRT1) activators were subjected to the integrated 2D and 3D QSAR approaches. The derived 3D QSAR models yielded high cross validated q^2^ values of 0.682 and 0.628 for CoMFA and CoMSIA respectively. The non-cross validated correlation values of r^2^_training_ = 0.89; predictive r^2^_test_ = 0.69 for CoMFA and r^2^=0.87; predictive r^2^_test_ =0.67 for CoMSIA reflected the statistical significance of the developed model. The steric, electrostatic, hydrophobic and hydrogen bond acceptor interactions have been found important in describing the variation in human SIRT1 activation. Further, 2D QSAR model for the same dataset yielded high statistical significance and derived 2D model’s parameters corroborated with 3D model in terms of features. The developed model was also validated through the available active conformation structure of SIRT1. Developed models may be useful for the identification of potential novel human SIRT1 activators as therapeutic agent.

## 1. Introduction

Human analogues of yeast Silent information regulator (SIR) protein also known as sirtuins (SIRT) belong to class III of histone de-acetylases family (HDACs) [1,2] These proteins are classified as SIRT 1-7 according to their sequence similarity and inherent requirement for nicotinamide adenine dinucleotide (NAD) in mammalian domain [3]. SIRTs are evolutionarily conserved enzymes that differ significantly from other classes of histone de-acetylases because of their specialized functions in metabolic pathways [4]. Mammalian SIRTs are known to affect numerous cellular functions related to aging, apoptosis, inflammation and endocrine signaling [5]. Seven types of mammalian sirtuins expressed in different tissues have been categorized in four classes of proteins: class I (SIRT1–3), class II (SIRT4), class III (SIRT5), and class IV (SIRT6–7) [3]. The therapeutic efficacy of human SIRT1 activation has been explored in several cardio-vascular diseases [6] modulation of mitochondrial biogenesis, metabolic rate, insulin sensitivity, glucose and lipid metabolism [7-11] and in several age-related diseases [15]. Cardiac specific over expression of SIRT1 has been known to prevent age-dependent increase in cardiac hypertrophy, apoptosis, cardiac dysfunction, and expression of senescence markers [6, 12]. SIRT’s catalytic domain consists two sub-domains; a large NAD+ binding subdomain and a small Zn2+-binding subdomain module has been resolved [13, 14]. SIRT1 activation is believed to be largely regulated by its allosteric site [15–16].

Resveratrol, a natural SIRT1 activator has been shown to improve the metabolic state of rodent models of obesity and type-2 diabetes mellitus [7, 17]. The SIRT1 activators could be mainly categorized into two classes: natural plant polyphenols and synthesized small molecules which include imidazo[1,2-b]thiazole derivatives [18], oxazolo[4,5-b]pyridines and their related heterocyclic analogs [19], 1,4-dihydropyridine analogs [20], resveratrol analogues [21, 22], and some quinoxaline analogs [23]. The crystal structure of human SIRT-1 protein with its small molecule activator was unknown until recently [24], thus few ligand-based studies have been reported on SIRT1 activators [25, 26]. However, no quantitative validation of the reported models has been reported in these studies. Our past successful efforts of integrated ligand- and structure-based drug discovery on other targets [27, 28, 29, 30] encouraged us to use this approach in the presented project. We herein report a ligand-based 3D QSAR models using known SIRT1 activators and its validation through available crystal structure of SIRT1.

## 2. Methodology

### 2.1. Dataset

The QSAR studies have been performed on a dataset of fifty compounds belonging to Imidazothiazole and Oxazolopyridine class having human SIRT1 activation potency with EC_1.5_ values ranging from 0.16μM to 130μM [18, 19]. Activation in the assay was measured as the concentration of compound required to increase the enzyme activity by 50% (EC_1.5_). The EC_1.5_ values were converted into negative logarithm of EC_1.5_ (pEC_1.5_) for the use in the QSAR studies. The fifty compounds in the dataset were rationally distributed into a training set of 26 molecules and a test set of 24 molecules to generate the QSAR models and evaluate the predictive ability of the developed models respectively. The chemical structures along with their biological activity of all the dataset compounds have been shown in Figure1, Table1 and Table 2.

**Figure 1.**
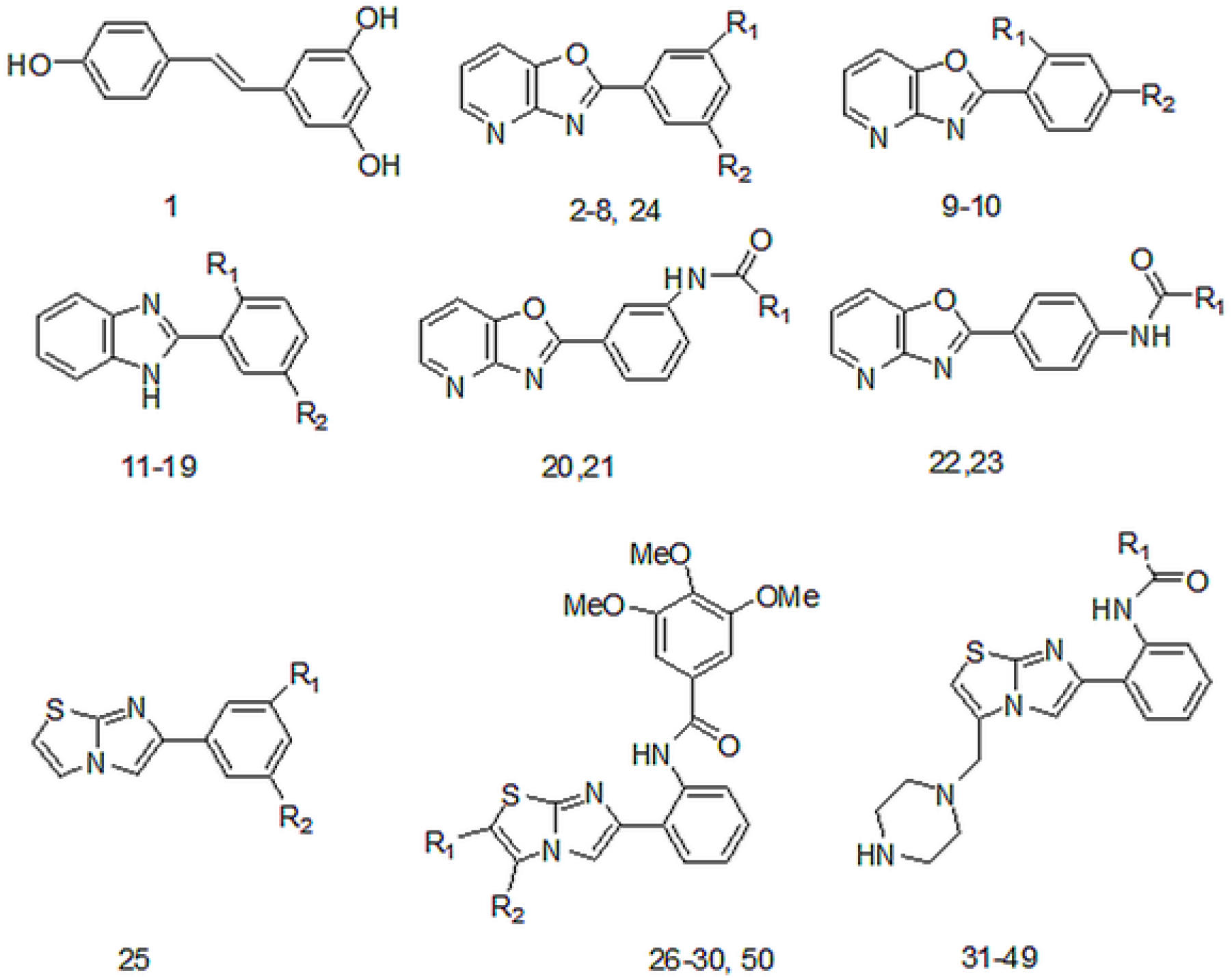
Common Structure of the SIRT1 activators used in the 3D-QSAR study.

**Table 1.**
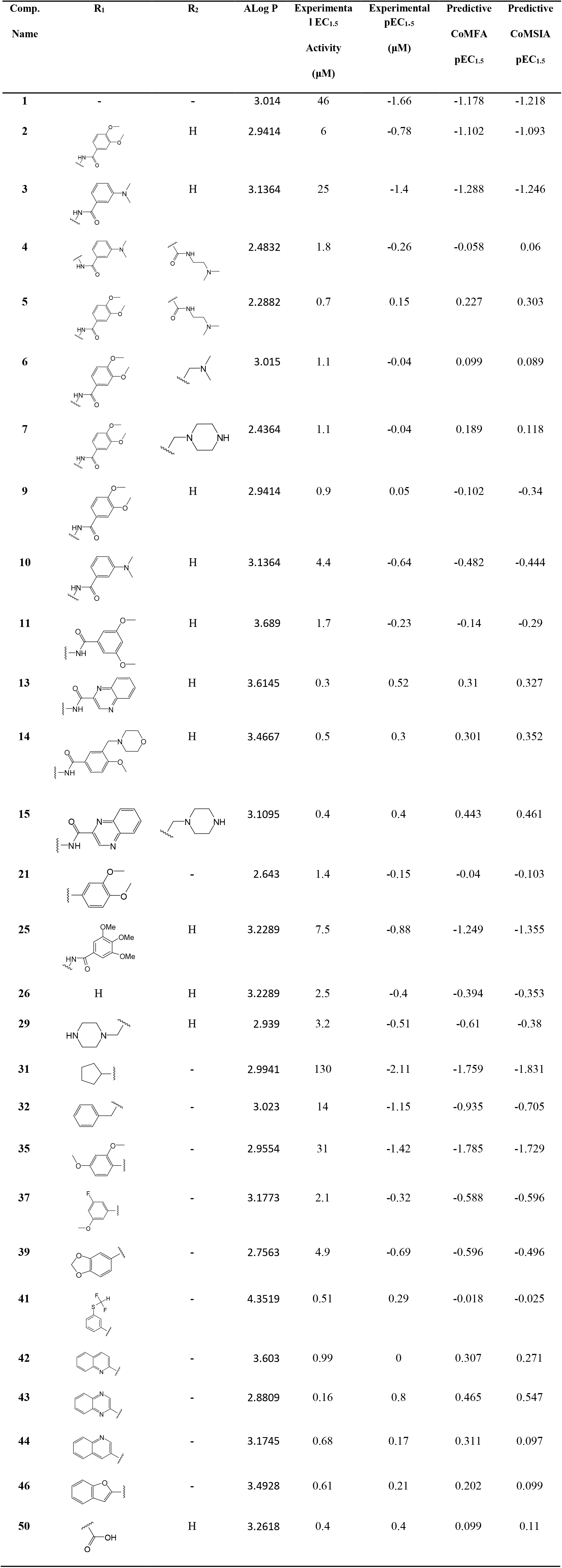
Structures of the training set compounds along with their experimental and predicted biological activity.

| Comp.<br>Name | R<sub>1</sub> | R<sub>2</sub> | ALog P | Experimental<br>1 EC<sub>1.5</sub><br>Activity<br>(μM) | Experimental<br>pEC<sub>1.5</sub><br>(μM) | Predictive<br>CoMFA<br>pEC<sub>1.5</sub> | Predictive<br>CoMSIA<br>pEC<sub>1.5</sub> |
| --- | --- | --- | --- | --- | --- | --- | --- |
| 1 | - | - | 3.014 | 46 | -1.66 | -1.178 | -1.218 |
| 2 |  | H | 2.9414 | 6 | -0.78 | -1.102 | -1.093 |
| 3 |  | H | 3.1364 | 25 | -1.4 | -1.288 | -1.246 |
| 4 |  |  | 2.4832 | 1.8 | -0.26 | -0.058 | 0.06 |
| 5 |  |  | 2.2882 | 0.7 | 0.15 | 0.227 | 0.303 |
| 6 |  |  | 3.015 | 1.1 | -0.04 | 0.099 | 0.089 |
| 7 |  |  | 2.4364 | 1.1 | -0.04 | 0.189 | 0.118 |
| 9 |  | H | 2.9414 | 0.9 | 0.05 | -0.102 | -0.34 |
| 10 |  | H | 3.1364 | 4.4 | -0.64 | -0.482 | -0.444 |
| 11 |  | H | 3.689 | 1.7 | -0.23 | -0.14 | -0.29 |
| 13 |  | H | 3.6145 | 0.3 | 0.52 | 0.31 | 0.327 |
| 14 |  | H | 3.4667 | 0.5 | 0.3 | 0.301 | 0.352 |
| 15 |  |  | 3.1095 | 0.4 | 0.4 | 0.443 | 0.461 |
| 21 |  | - | 2.643 | 1.4 | -0.15 | -0.04 | -0.103 |
| 25 |  | H | 3.2289 | 7.5 | -0.88 | -1.249 | -1.355 |
| 26 | H | H | 3.2289 | 2.5 | -0.4 | -0.394 | -0.353 |
| 29 |  | H | 2.939 | 3.2 | -0.51 | -0.61 | -0.38 |
| 31 |  | - | 2.9941 | 130 | -2.11 | -1.759 | -1.831 |
| 32 |  | - | 3.023 | 14 | -1.15 | -0.935 | -0.705 |
| 35 |  | - | 2.9554 | 31 | -1.42 | -1.785 | -1.729 |
| 37 |  | - | 3.1773 | 2.1 | -0.32 | -0.588 | -0.596 |
| 39 |  | - | 2.7563 | 4.9 | -0.69 | -0.596 | -0.496 |
| 41 |  | - | 4.3519 | 0.51 | 0.29 | -0.018 | -0.025 |
| 42 |  | - | 3.603 | 0.99 | 0 | 0.307 | 0.271 |
| **43** |  | - | 2.8809 | 0.16 | 0.8 | 0.465 | 0.547 |
| **44** |  | - | 3.1745 | 0.68 | 0.17 | 0.311 | 0.097 |
| **46** |  | - | 3.4928 | 0.61 | 0.21 | 0.202 | 0.099 |
| **50** |  | H | 3.2618 | 0.4 | 0.4 | 0.099 | 0.11 |

**Table 2.**
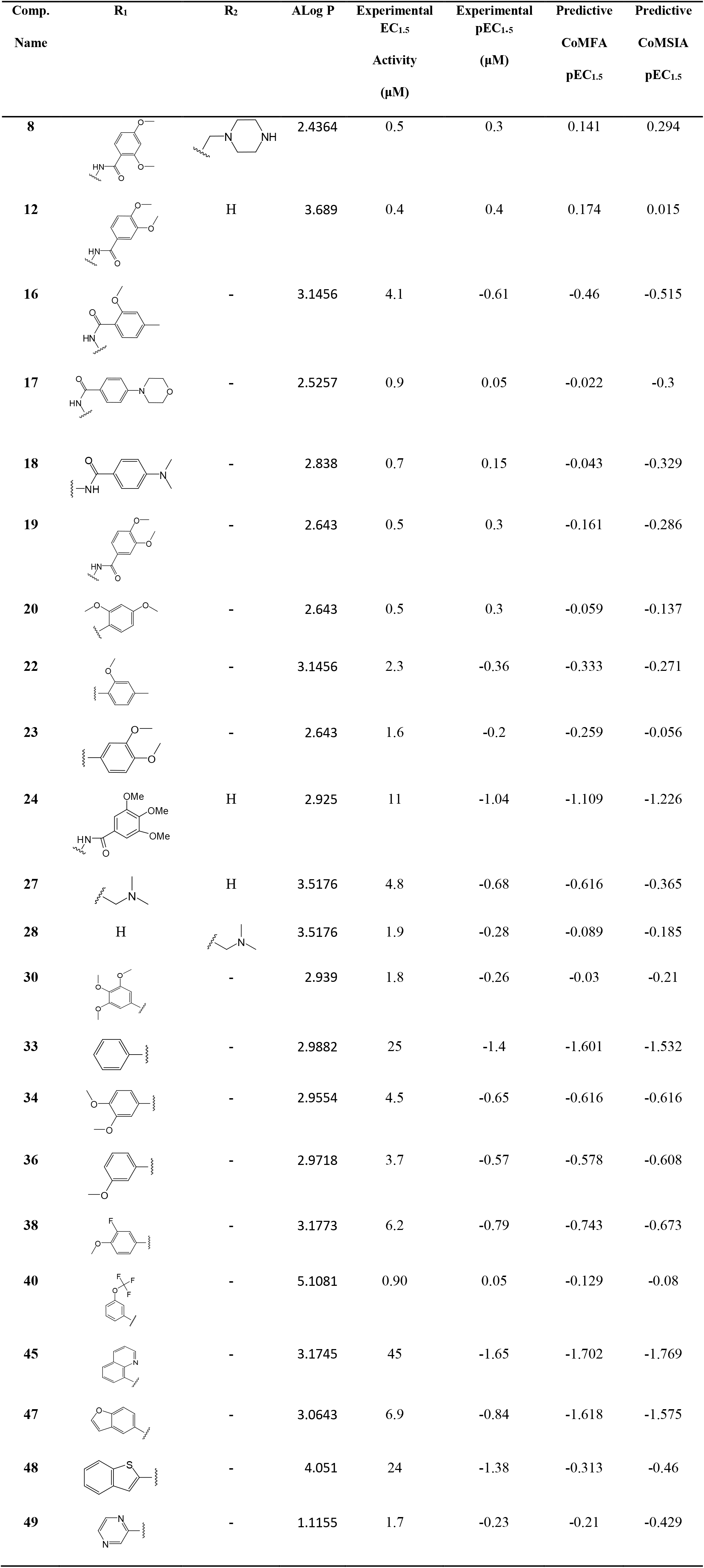
Structures of the test set compounds along with their experimental and predicted biological activity.

| Comp.<br>Name | R<sub>1</sub> | R<sub>2</sub> | ALog P | Experimental<br>EC<sub>1.5</sub><br>Activity<br>(μM) | Experimental<br>pEC<sub>1.5</sub><br>(μM) | Predictive<br>CoMFA<br>pEC<sub>1.5</sub> | Predictive<br>CoMSIA<br>pEC<sub>1.5</sub> |
| --- | --- | --- | --- | --- | --- | --- | --- |
| 8 |  |  | 2.4364 | 0.5 | 0.3 | 0.141 | 0.294 |
| 12 |  | H | 3.689 | 0.4 | 0.4 | 0.174 | 0.015 |
| 16 |  | - | 3.1456 | 4.1 | -0.61 | -0.46 | -0.515 |
| 17 |  | - | 2.5257 | 0.9 | 0.05 | -0.022 | -0.3 |
| 18 |  | - | 2.838 | 0.7 | 0.15 | -0.043 | -0.329 |
| 19 |  | - | 2.643 | 0.5 | 0.3 | -0.161 | -0.286 |
| 20 |  | - | 2.643 | 0.5 | 0.3 | -0.059 | -0.137 |
| 22 |  | - | 3.1456 | 2.3 | -0.36 | -0.333 | -0.271 |
| 23 |  | - | 2.643 | 1.6 | -0.2 | -0.259 | -0.056 |
| 24 |  | H | 2.925 | 11 | -1.04 | -1.109 | -1.226 |
| 27 |  | H | 3.5176 | 4.8 | -0.68 | -0.616 | -0.365 |
| 28 | H |  | 3.5176 | 1.9 | -0.28 | -0.089 | -0.185 |
| 30 |  | - | 2.939 | 1.8 | -0.26 | -0.03 | -0.21 |
| 33 |  | - | 2.9882 | 25 | -1.4 | -1.601 | -1.532 |
| 34 |  | - | 2.9554 | 4.5 | -0.65 | -0.616 | -0.616 |
| 36 |  | - | 2.9718 | 3.7 | -0.57 | -0.578 | -0.608 |
| 38 |  | - | 3.1773 | 6.2 | -0.79 | -0.743 | -0.673 |
| 40 |  | - | 5.1081 | 0.90 | 0.05 | -0.129 | -0.08 |
| 45 |  | - | 3.1745 | 45 | -1.65 | -1.702 | -1.769 |
| 47 |  | - | 3.0643 | 6.9 | -0.84 | -1.618 | -1.575 |
| 48 |  | - | 4.051 | 24 | -1.38 | -0.313 | -0.46 |
| 49 |  | - | 1.1155 | 1.7 | -0.23 | -0.21 | -0.429 |

### 2.2. Computational approach and Molecular alignment

The 3D-QSAR techniques such as CoMFA and CoMSIA have been widely used in computer-aided drug discovery to understand the physicochemical features required for drug–receptor interaction [31-32]. In present study 3D-QSAR studies were performed on an SGI Octane R12000 workstation using SYBYL 6.9 molecular modeling software [33]. Structural sketches of all the SIRT1 activators were drawn in SYBYL. Partial charges of these ligands were calculated using Gasteiger-Hückel method and their geometry were optimized using Tripos force field using standard SYBYL settings. “Multi-search” method of conformational search was performed using the following settings: maximum cycles (400), maximum conformers (400), tolerance (0.40), energy cutoff (70 kcal/mol), rms gradient (3.0) and number of hit (12). The lowest energy conformation obtained for each molecule were used in the analysis. The core (shown in bold) of the most active compound **43** (Figure 2A) was used as a template for molecular alignment and the calculated pEC_1.5_ values were used as dependent variables for carrying out PLS analysis in CoMFA and CoMSIA study. The overall alignment of the training set molecules has been shown in Figure2B.

**Figure 2.**
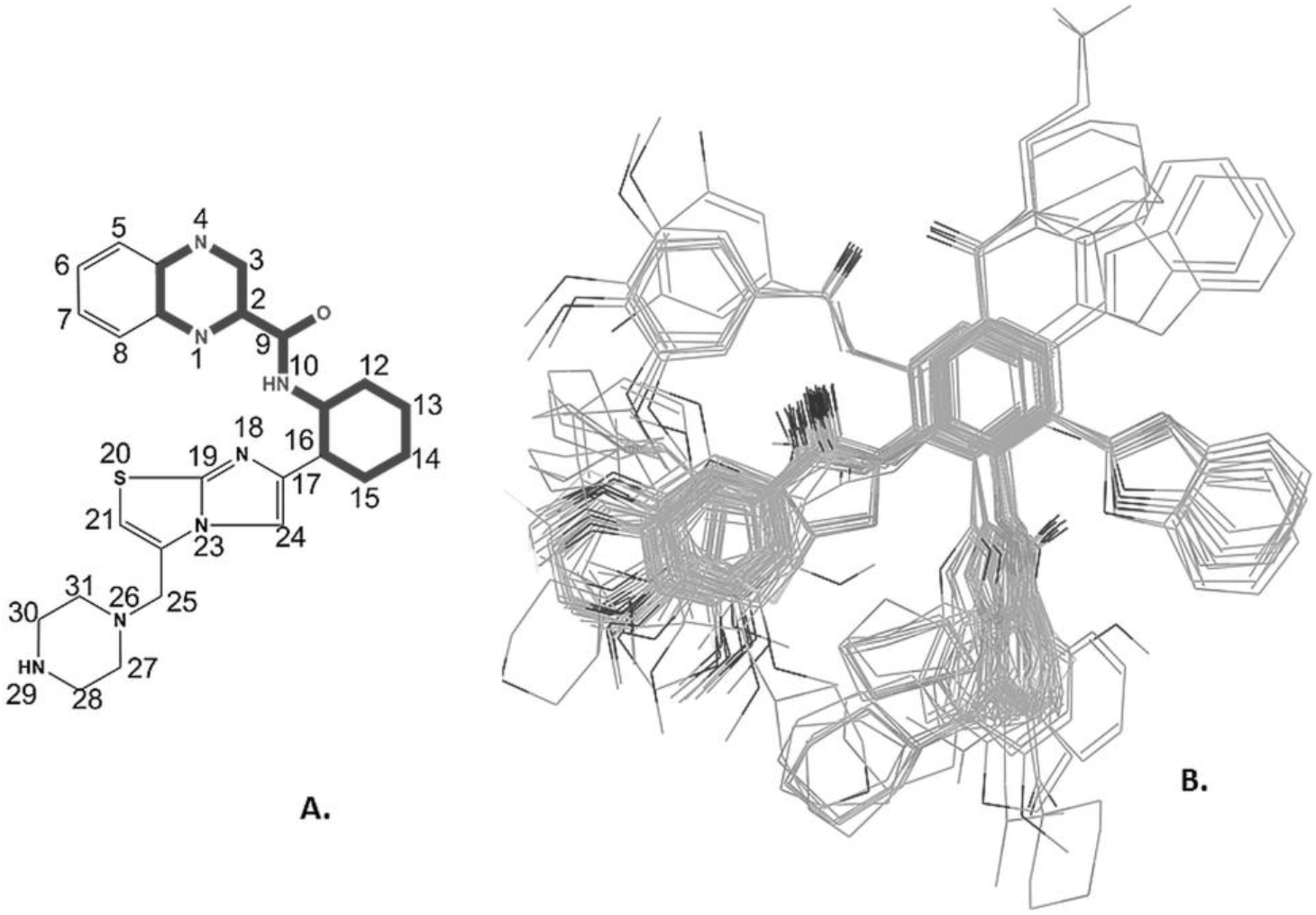
**(A)** The core structure (shown in bold) of the most active molecule of the dataset (compound **43**) and the numbering system used in the 3D-QSAR study. **(B)** The overall alignment of the training set molecules used in the 3D-QSAR study.

### 2.3. CoMFA and CoMSIA model generation

#### 2.3.1. CoMFA

Aligned molecules of the training set were aggregated in a 3D cubic lattice with a grid spacing of 2.0 Å in the x, y and z directions for generating the CoMFA fields. Steric (Lennard–Jones potential) and electrostatic (Columbic) potential energies were calculated at each lattice point using Tripos force field and a distance dependent dielectric constant of 1.0. A sp^3^-hybridised carbon with a charge of +1.0 and a Van der Waals radius of 1.52 Å was used as a probe to calculate various steric and electrostatic fields. An energy cutoff value of 30 kcal/mol was applied to all CoMFA calculations to avoid excessively high and unrealistic energy values within the molecule.

#### 2.3.2. CoMSIA

CoMSIA is based on the molecular similarity indices in the same lattice box used for the CoMFA calculations [29]. During CoMFA steric and electrostatic calculation, several grid points on the molecular surface can be neglected because of the rapid increase in Van der Waals repulsion. CoMSIA uses a Gaussian-type function based on distance to avoid a dramatic change in the potential energy of the grid points near the molecular surface. Thus, CoMSIA is considered superior to CoMFA technique and can obtain more stable models than CoMFA. In the present study, steric, electrostatic, hydrophobic, H-donor and H-acceptor similarity fields were calculated using the default CoMSIA settings (Probe with charge +1, radius 1 Å, grid spacing 2 Å, attenuation factor of 0.3 and hydrophobic, hydrogen-bond doner, and hydrogen-bond acceptor as +1).

#### 2.3.3. Partial least squares (PLS) and Predictive r_2_ analysis

SIRT1 activity was correlated with the coefficient of steric, electrostatic and hydrophobic potentials obtained from CoMFA and CoMSIA. Leave one out (LOO) validation was utilized as a measure to rule out chances of false correlation. To further assess the statistical robustness of the derived models, a group cross validation using 20 groups, repeating the procedure 20 times was also carried out. The mean of 20 readings is given as r^2^_cv(mean)_. The full PLS analysis was carried out with a column filtering of 2.0 kcal/mol. The predictive r^2^ value is based on test set molecules and defined as

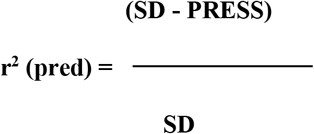

where SD is the sum of squared deviation of each biological activity from the mean, and PRESS is the sum of squared deviations between the actual and the predicted activities [30].

### 2.4. 2D QSAR model generation

In *2D QSAR* analysis constitutional, topological descriptors, BCUT, geometrical descriptors were calculated using PaDEL-Descriptor software [34]. Linear regression was performed using Systat (v12.2) software using the SIRT1 pEC_1.5_ as dependent and ligand-based physicochemical parameters as independent parameters. Pearson correlation matrix was constructed to determine the inter-correlation between the independent parameters and the correlation between the dependent and independent parameters. Stepwise multi-parameter regression analysis was performed by backward elimination method using Systat software (v12.2). These descriptors were then used to generate an equation which would give insight into the various physicochemical and structural parameters influencing SIRT1 activation.

### 2.5. Molecular docking

The SIRT1 protein structure (PDB ID: 4ZZH) was prepared for docking using the protein preparation wizard of the Schrodinger Suite of programs (Small molecule drug discovery suite2015-2, S., LLC, NY. 2015) with default options to assign bond order and remove water molecules that are 5 Å away from hetero groups. H-bond assignment was done using PROPKA at pH 7.0 and the orientation of hydroxyl groups, amide groups of Asn and Gln, and charge state of His residues were optimized. At the end, protein was minimized to an RMSD limit from the starting structure of 0.3 Å with the OPLS3 force field. Binding site/docking grid was defined by selecting co-crystallized small molecule SIRT1 activator. The ligand libraries were prepared using LigPrep with Epik at pH 7.0 while retaining specific chirality of the ligands. Partial atomic charges were computed using the OPLS-3 force field and lowest energy conformation was retained. The lowest energy conformation among the largest populated cluster sharing common interactions was selected as the best binding pose for docked ligands.

## 3. Results and discussion

CoMFA and CoMSIA methods were used to generate 3D-QSAR predictive models on a set of 50 chemically diverse human SIRT1 activators. Best performance 3D-QSAR model was selected based on the statistical parameters obtained among the various 3D-QSAR models generated from the training set and its predictive ability to discriminate inactive and active molecules using test set compounds. The results of the CoMFA, CoMSIA studies have been summarized in Table 3.

**Table. 3.** PLS statistics of CoMFA (TS) and CoMSIA (SEHA) models.

| Parameters |  | CoMFA (TS) | CoMSIA (SEHA) |
| --- | --- | --- | --- |
| $q^2$ | | 0.682 | 0.628 |
| PRESS |  | 0.427 | 0.455 |
| $r^2$ | | 0.889 | 0.870 |
| SEE |  | 0.247 | 0.267 |
| F |  | 100.557 | 83.939 |
| N |  | 2 | 2 |
| Fractions | S | 0.384 | 0.164 |
|  | E | 0.616 | 0.320 |
|  | H | - | 0.241 |
|  | A | - | 0.276 |
| $r^2_{bs}$ (20runs) | | 0.931 | 0.917 |
| SD<sub>bs</sub> |  | 0.029 | 0.020 |
| $r^2_{CV (mean)}$ (20runs) | | 0.670 | 0.625 |
| $r^2_{pred}$ | | 0.694 | 0.666 |

$q^2$ = leave one out cross-validation correlation coefficient; PRESS = LOO cross-validated standard error / predictive sum of squares; $r^2$ = conventional correlation; SEE = standard error of estimate; F = degree of freedom; N = optimal number of component; $r^2_{bs}$ = bootstrapping correlation; SD<sub>bs</sub> = bootstrapping standard deviation; $r^2_{CV (mean)}$ = group cross-validation; TS = Tripos standard; SEHA = Steric, electrostatic, Hydrophobic and Acceptor.

### 3.1. CoMFA analysis

The CoMFA model with cross validated q^2^ of 0.682 for 2 principal components and a non-cross validated conventional r^2^ of 0.889, F value of 100.56 and standard error of estimate (SEE) of 0.247 was selected for further studies. Bootstrapping analysis (20 runs) result, r^2^_bs_ of 0.931 (SD_bs_ = 0.029) established the robustness of this model. In addition to LOO, a group cross validation was further done to assess the internal predictive ability of the model. The mean r^2^_CV_ of 0.670 (TS) value obtained from the 20 times cross-validation with 20 groups suggests model’s excellent internal predictability (Table 3). The external predictive r^2^ of 0.694 on a test set of 24 SIRT1 activators showed good predictive ability of the generated model (Figure 3A). The predictive pEC_1.5_ values of the training as well as test set molecules based on the CoMFA model have been included in the Table 1 and Table 2 respectively.

**Figure 3.**
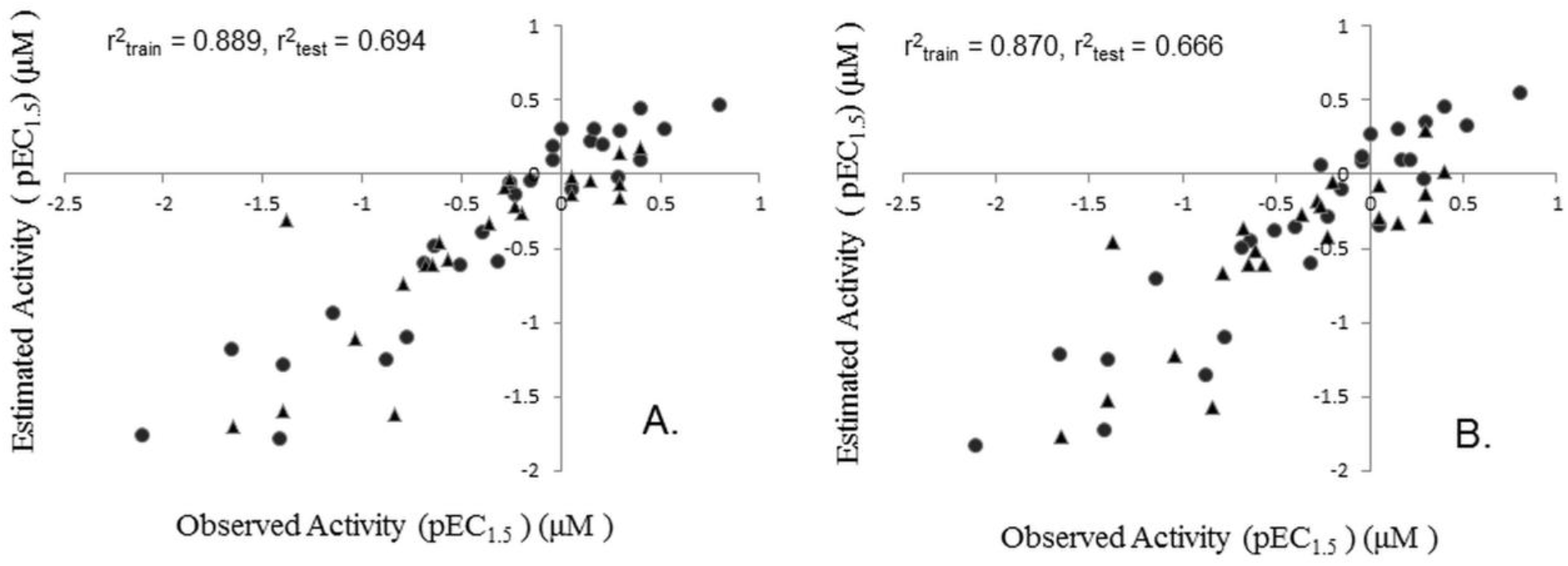
Correlation graph between observed and predicted activities of training set (dots) and test set (triangles) molecules **(A)** CoMFA and **(B)** CoMSIA.

### 3.2. CoMSIA analysis

The CoMSIA model having steric(S), electrostatic (E), hydrophobic (H) and hydrogen bond acceptor (A) descriptors gave the highest q^2^ value of 0.628 at 2 components and a conventional non cross-validated r^2^ of 0.870, F value of 83.94 and standard error of estimate SEE of 0.267. Bootstrapping analysis (20 runs) value, r^2^_bs_ of 0.917 with low standard deviation of 0.020 suggests statistical ability and robustness of the model. Like CoMFA, the mean r^2^_cv_ value of 0.625 with 20 groups for CoMSIA model revealed that the model has high internal predictivity. The predictive r^2^ of 0.666 for the 24 test set compounds showed the usefulness of the model in external predictivity (Figure 3B). The Predictive pEC_1.5_ values of the training as well as test set molecules based on the CoMSIA model have been included in the Table1 and Table 2 respectively.

### 3.3. CoMFA and CoMSIA contour maps analysis

3D contour maps were generated from the coefficients of CoMFA and CoMSIA models. These contours represent physicochemical regions in 3D space around the dataset molecules which are responsible in determining the activity and explores the importance of various substituents in their 3D orientation. The Contour maps generated from the CoMFA and CoMSIA models are shown in Figure 4.

**Figure 4.**
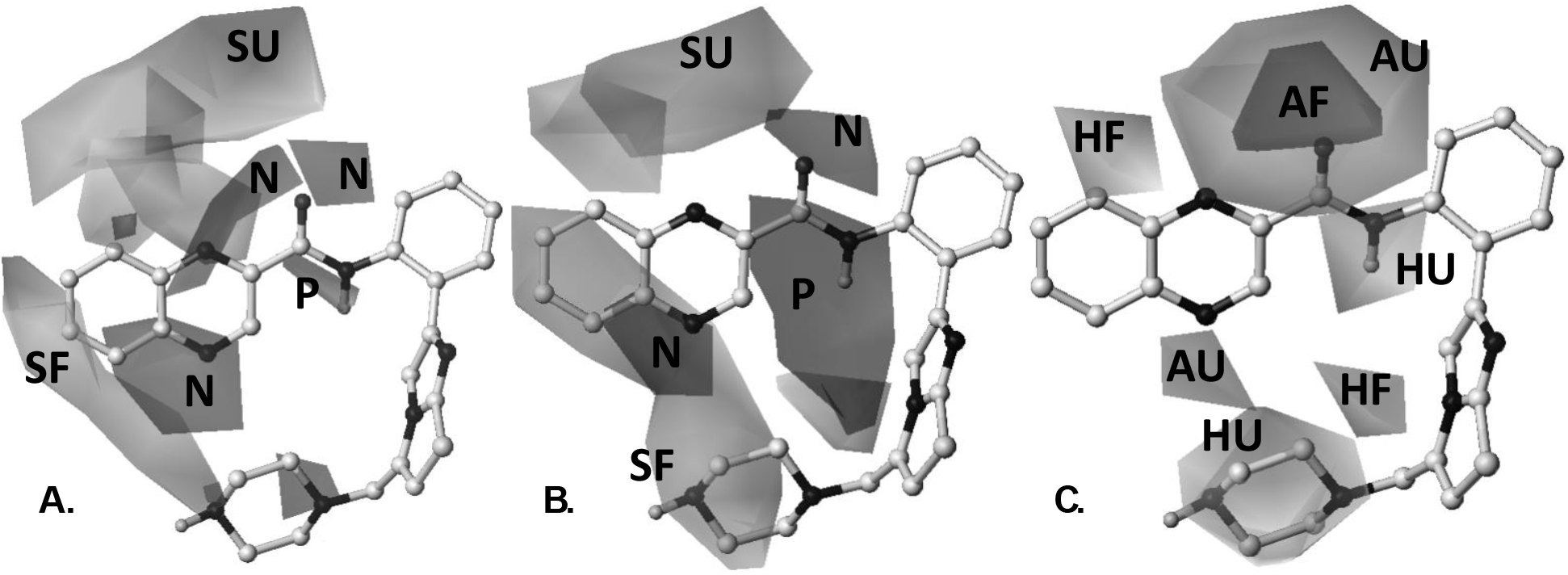
PLS contours from 3D-QSAR analysis of SIRT1 activators **(A)** Steric and electrostatic CoMFA contours; **(B)** Steric and electrostatic CoMSIA contours. The ‘SF’ and ‘SU’ labeled contours indicate sterically favored and disfavored regions respectively while the ‘P’ and ‘N’ labeled contours denote regions that favor electropositive substituent and electronegative substituent respectively; **(C)** hydrogen bond acceptor and hydrophobic contours. The ‘AF’ and ‘AU’ labeled polyhedral represents the favored and disfavored regions respectively for hydrogen bond acceptor feature. Similarly, ‘HF’ and ‘HU’ contours represents hydrophobically favored and dis favored regions.

An analysis of contrast of CoMFA and CoMSIA contours around compound **43 (**most active molecule of the dataset) signify the importance of sterically favorable region (green color) near the 6, 7, 8 position of quinoxaline moiety for SIRT1 activation. The higher activities among all the active compounds of the training set (viz. compound **43, 13, 50, 15, 14, 27, 41, 46, 44, 5** and **9**) can be explained based on presence of aromatic ring of the quinoxaline moiety at the same position. It has been also found that an addition of bulky group near 29, and 30 positions of the piperazine moiety further increased the SIRT1 activation potency (*viz.* compound **43, 14, 41, 46, 44, 5** and **42**). The absence of such functionality in 3D space led to the decrease in the activation potency (viz. compound **2, 25, 32, 3, 35, 1**, and **31**). The sterically unfavorable region at the upper side of the 4, 5 positions of quinoxaline ring as well as on the lower plane of the piperazine moiety has been found important for SIRT 1 activation (Figure 4A).

When the electrostatic contours were considered, both the CoMFA and CoMSIA contours emphasized the importance of the electropositive substituent near the amide moeity and the presence of substituent with high electronegative functionality in the region adjoining the nitrogen atom at the 9 positions to the keto oxygen of the quinoxaline ring, for high SIRT1 activation potency. These contours also conclude the fact that optimal activation potency is achieved when the electronegative functionality at that region is being contributed by the nitrogen atom (Figure 4A and 4B).

The PLS contour analysis of the hydrogen bond acceptor feature highlighted the importance of the presence of a hydrogen bond acceptor atom in the vicinity of the ketonic oxygen atom at position 9 of the molecule. These contours well explain the variation in biological activity among the compounds of series (compounds **1, 2, 3, 25, 31**, and **35** failed to map the H-bond acceptor group at this 3D space represent least potent molecules of this series). The space present in between and behind the phenyl ring and the piperazine ring has been found unfavorable for acceptor functionality (Figure 4C). In the case of hydrophobic contours, the hydrophobic favorable regions near the 5 position of quinoxaline ring and around piperazine nucleus while hydrophobic disfavored regions around the CONH functionality of the molecule **43** facilitate the effective binding of SIRT1 activators to the target protein (Figure 4C).

### 3.4. 2D QSAR Analysis

CoMFA and CoMSIA predictive models can be sensitive to ligand alignment [35]. Therefore, to rule out any false positive results, and validate the model we performed alignment free 2D QSAR analysis for the same dataset. Different combination of physiochemical parameters showing statistical correlation with the biological activity were carried out using stepwise multiple regression analysis to develop meaningful QSAR equations.

We derived structure activity relationship equation for all the 50 dataset ligands. The generated equation showed the importance of Polarizability (AMR) [36], BCUT (SPMAX3_BHS) [37] and topological parameter (WPOL) [38] in defining the structure activity relationship of SIRT1 activators. Regression analysis of the above physico-chemical parameters with biological activity generated following equation:

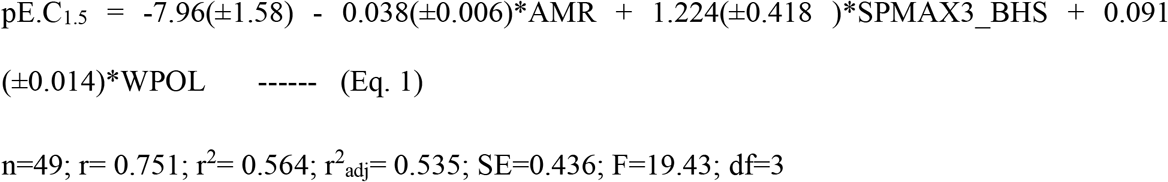

Eq. 1 showed 99.99% significance with the correlation coefficient value of 0.75 between the predicted and observed activity (Figure 5) after removal of one outlier compound (Compound **45**). The low standard error of estimate(s) and a high F value suggests that the model is highly statistically significant. Values of physico-chemical parameters used in the equations as well as observed and predicted activities of all compounds in training and test set listed in Table 4.

**Figure 5.**
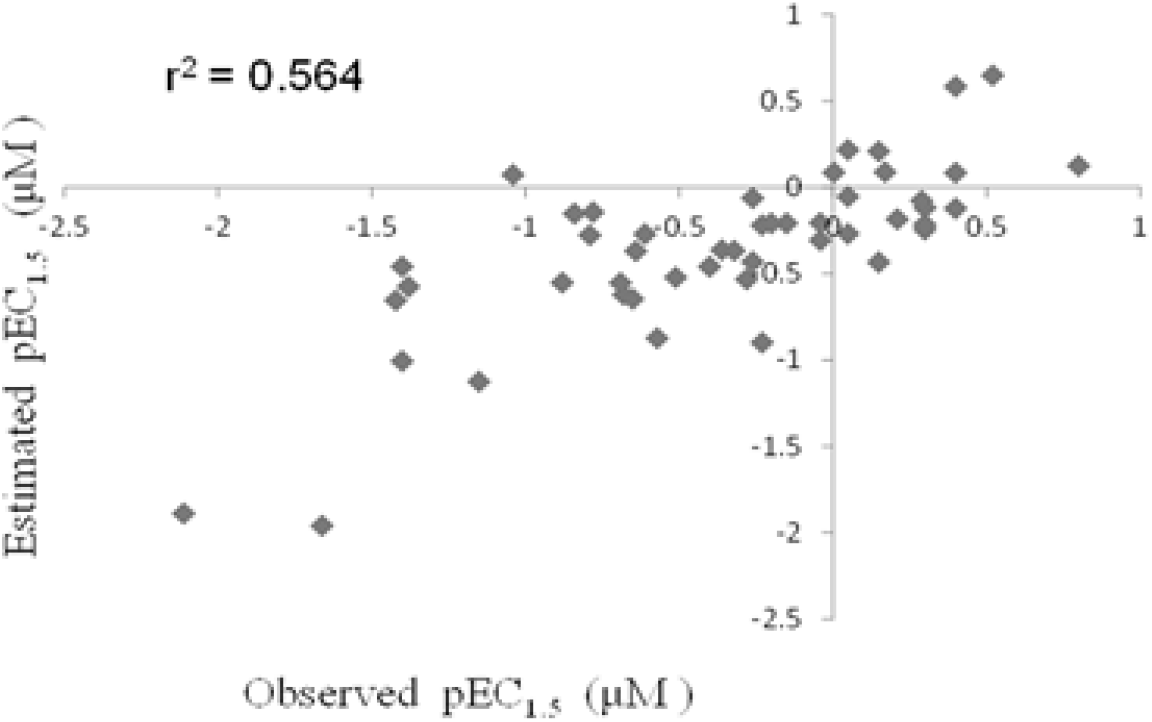
2D Correlation graph between observed and predicted activities of dataset set ligands.

**Table 4.**
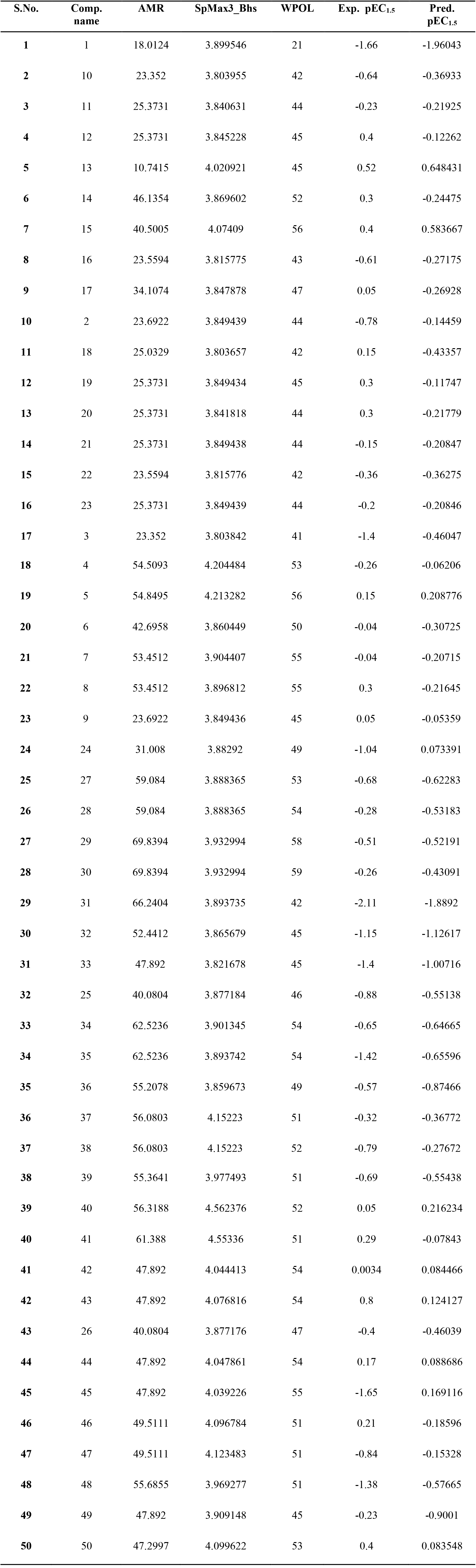
Values of descriptors used in the 2D QSAR equation as well as observed and predicted activities of all compounds in data set.

| S.No. | Comp. name | AMR | SpMax3_Bhs | WPOL | Exp. pEC<sub>1.5</sub> | Pred. pEC<sub>1.5</sub> |
| --- | --- | --- | --- | --- | --- | --- |
| 1 | 1 | 18.0124 | 3.899546 | 21 | -1.66 | -1.96043 |
| 2 | 10 | 23.352 | 3.803955 | 42 | -0.64 | -0.36933 |
| 3 | 11 | 25.3731 | 3.840631 | 44 | -0.23 | -0.21925 |
| 4 | 12 | 25.3731 | 3.845228 | 45 | 0.4 | -0.12262 |
| 5 | 13 | 10.7415 | 4.020921 | 45 | 0.52 | 0.648431 |
| 6 | 14 | 46.1354 | 3.869602 | 52 | 0.3 | -0.24475 |
| 7 | 15 | 40.5005 | 4.07409 | 56 | 0.4 | 0.583667 |
| 8 | 16 | 23.5594 | 3.815775 | 43 | -0.61 | -0.27175 |
| 9 | 17 | 34.1074 | 3.847878 | 47 | 0.05 | -0.26928 |
| 10 | 2 | 23.6922 | 3.849439 | 44 | -0.78 | -0.14459 |
| 11 | 18 | 25.0329 | 3.803657 | 42 | 0.15 | -0.43357 |
| 12 | 19 | 25.3731 | 3.849434 | 45 | 0.3 | -0.11747 |
| 13 | 20 | 25.3731 | 3.841818 | 44 | 0.3 | -0.21779 |
| 14 | 21 | 25.3731 | 3.849438 | 44 | -0.15 | -0.20847 |
| 15 | 22 | 23.5594 | 3.815776 | 42 | -0.36 | -0.36275 |
| 16 | 23 | 25.3731 | 3.849439 | 44 | -0.2 | -0.20846 |
| 17 | 3 | 23.352 | 3.803842 | 41 | -1.4 | -0.46047 |
| **18** | 4 | 54.5093 | 4.204484 | 53 | -0.26 | -0.06206 |
| **19** | 5 | 54.8495 | 4.213282 | 56 | 0.15 | 0.208776 |
| **20** | 6 | 42.6958 | 3.860449 | 50 | -0.04 | -0.30725 |
| **21** | 7 | 53.4512 | 3.904407 | 55 | -0.04 | -0.20715 |
| **22** | 8 | 53.4512 | 3.896812 | 55 | 0.3 | -0.21645 |
| **23** | 9 | 23.6922 | 3.849436 | 45 | 0.05 | -0.05359 |
| **24** | 24 | 31.008 | 3.88292 | 49 | -1.04 | 0.073391 |
| **25** | 27 | 59.084 | 3.888365 | 53 | -0.68 | -0.62283 |
| **26** | 28 | 59.084 | 3.888365 | 54 | -0.28 | -0.53183 |
| **27** | 29 | 69.8394 | 3.932994 | 58 | -0.51 | -0.52191 |
| **28** | 30 | 69.8394 | 3.932994 | 59 | -0.26 | -0.43091 |
| **29** | 31 | 66.2404 | 3.893735 | 42 | -2.11 | -1.8892 |
| **30** | 32 | 52.4412 | 3.865679 | 45 | -1.15 | -1.12617 |
| **31** | 33 | 47.892 | 3.821678 | 45 | -1.4 | -1.00716 |
| **32** | 25 | 40.0804 | 3.877184 | 46 | -0.88 | -0.55138 |
| **33** | 34 | 62.5236 | 3.901345 | 54 | -0.65 | -0.64665 |
| **34** | 35 | 62.5236 | 3.893742 | 54 | -1.42 | -0.65596 |
| **35** | 36 | 55.2078 | 3.859673 | 49 | -0.57 | -0.87466 |
| **36** | 37 | 56.0803 | 4.15223 | 51 | -0.32 | -0.36772 |
| **37** | 38 | 56.0803 | 4.15223 | 52 | -0.79 | -0.27672 |
| **38** | 39 | 55.3641 | 3.977493 | 51 | -0.69 | -0.55438 |
| **39** | 40 | 56.3188 | 4.562376 | 52 | 0.05 | 0.216234 |
| **40** | 41 | 61.388 | 4.55336 | 51 | 0.29 | -0.07843 |
| **41** | 42 | 47.892 | 4.044413 | 54 | 0.0034 | 0.084466 |
| **42** | 43 | 47.892 | 4.076816 | 54 | 0.8 | 0.124127 |
| **43** | 26 | 40.0804 | 3.877176 | 47 | -0.4 | -0.46039 |
| **44** | 44 | 47.892 | 4.047861 | 54 | 0.17 | 0.088686 |
| **45** | 45 | 47.892 | 4.039226 | 55 | -1.65 | 0.169116 |
| **46** | 46 | 49.5111 | 4.096784 | 51 | 0.21 | -0.18596 |
| **47** | 47 | 49.5111 | 4.123483 | 51 | -0.84 | -0.15328 |
| **48** | 48 | 55.6855 | 3.969277 | 51 | -1.38 | -0.57665 |
| **49** | 49 | 47.892 | 3.909148 | 45 | -0.23 | -0.9001 |
| **50** | 50 | 47.2997 | 4.099622 | 53 | 0.4 | 0.083548 |

Table 5 lists the Pearson Correlation Matrix of the significant parameters used in the equation.

**Table 5:** Pearson Correlation Matrix of the significant parameters used in the equation.

|  | **AMR** | **SPMAX3_BHS** | **WPOL** |
| --- | --- | --- | --- |
| **AMR** | 1.000 |  |  |
| **SPMAX3_BHS** | 0.431 | 1 |  |
| **WPOL** | 0.707 | 0.431 | 1.000 |

The correlation between observed and predicted activities of all compounds using Eq. 1 has been represented graphically as shown in Figure 5.

Significant contribution of lipophilicity (AMR) parameter suggests the role of hydrophobic and van-der-Waals interactions in defining the variation in the biological activity of SIRT1 activators. Further, contribution of BCUT (SPMAX3_BHS) and topological parameter (WPOL) suggests the importance of hydrogen bonds, charge transfer interactions and steric hindrance geometric fit for SIRT-1 activation. These features are consistent with our 3D CoMFA and CoMSIA model where importance of all these features has been well explained.

### 3.5. Structure based validation of CoMFA, CoMSIA model

A recently reported structure of SIRT1-enzyme-activator complex [24] enabled us to validate our CoMFA, CoMSIA model. Figure 6 represents lowest energy conformation of most active compound bound with active form of SIRT1. Quinoxaline moiety of compound **43** has made hydrophobic contact with Ile227 and Ile223 at its 4, 5, 6 position which corroborated with hydrophobically favored cotutors of CoMFA and CoMSIA. Interaction of small molecule with Glu230 is very critical for SIRT1 activation. Glu230 also plays pivotal role of an electrostatic interaction between Glu230 and Arg446 in the activated conformation. Presence of any bulkier group will negatively impact the SIRT1 activation as shown in hydrophobic unfavorable contours of Figure 4. The Nitrogen atom of piperazine ring of compound **43** makes the H-bond interaction with Glu230 (Figure 6) well captured by electronegative contours in Figure 4. Other H-bond interactions between amide bond of compound **43** and residue Asn226, and Ile223 were well predicted by H-bond contours of CoMSIA model. In summary, our CoMFA, CoMSIA models predict the binding site interactions of small molecule SIRT1 activators very well.

**Figure 6.**
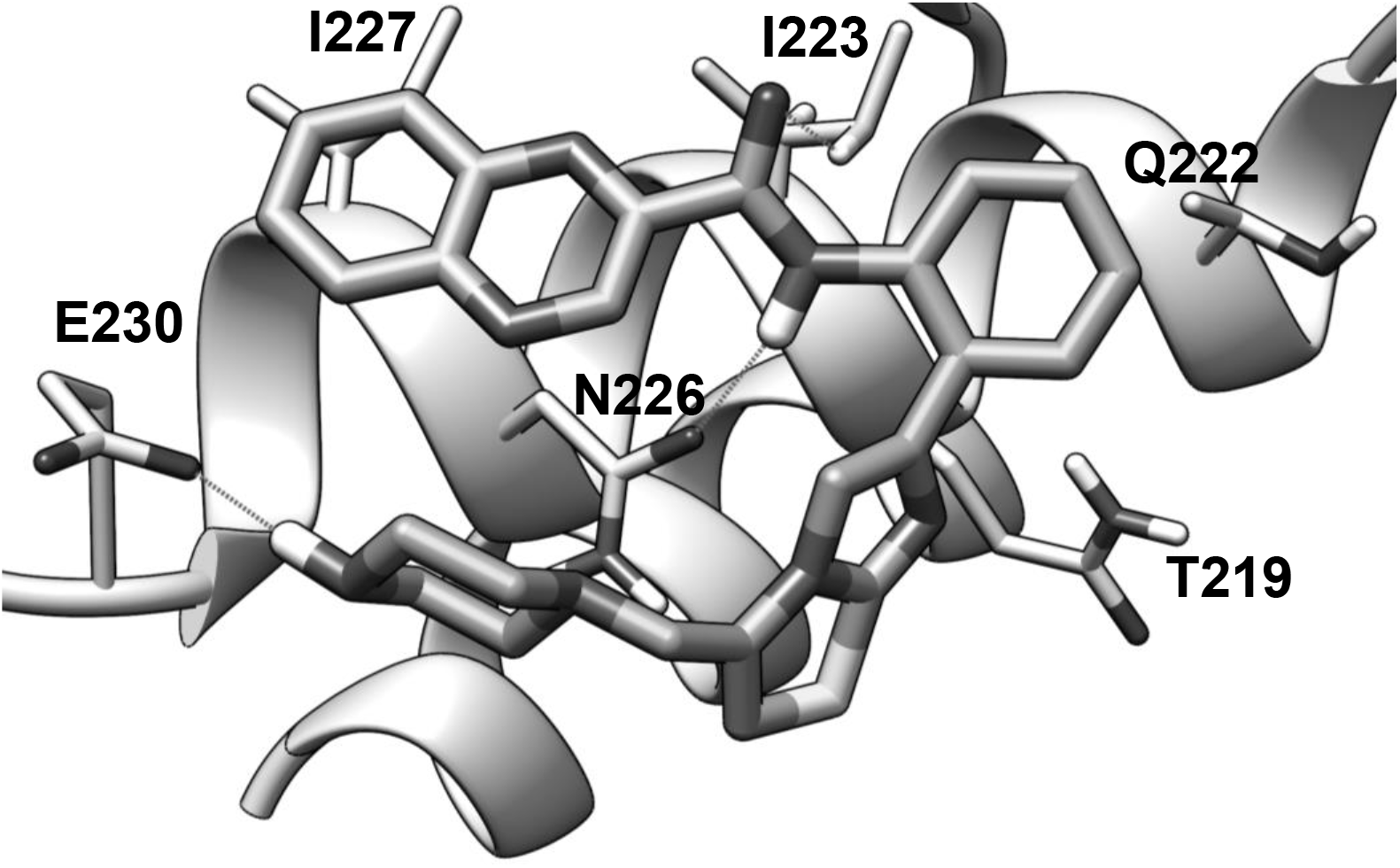
Lowest energy conformation of most active compound **43** bound with active form of SIRT1 protein.

## 4. Conclusion

The CoMFA and CoMSIA have been applied successfully to rationalize the structurally diverse SIRT1 activators. Steric, electrostatic, hydrophobic and hydrogen bond acceptor interactions in 3D space have been found important for SIRT1 activation. The developed 3D models showed excellent statistical significance in internal validation (q^2^, group cross-validation and bootstrapping) and performed very well in predicting the SIRT1 activation by the compounds in the test set. The developed model was also validated by 2D QSAR model as well as the crystal structure of active conformation of SIRT1. These models alone or in combination may be useful in the designing novel human SIRT1 activators with increased potency.

## Acknowledgement

This work was supported by the Mid-career Researcher Program (NRF-2017R1A2B4010084), Bio & Medical Technology Development Program (No. 2018M3A9A7057263), and Medical Research Center (MRC) grant (No. 2018R1A5A2025286) funded by the Ministry of Science and ICT (MSIT) through the National Research Foundation of Korea (NRF).

The authors declare no conflict of interests.

## References

[1] Blander, G.; Guarente, L. The Sir2 family of protein deacetylases. Annu. Rev. Biochem, 2004, 73, 417–435.

[2] Khochbin, S.; Verdel, A.; Lemercier, C.; Seigneurin-Berny, D. Functional significance of histone deacetylase diversity. Curr. Opin. Genet. Dev. 2001, 11, 162–166.

[3] Schwer, B.; Verdin, E. Conserved metabolic regulatory functions of sirtuins. Cell Metab. 2008, 7, 104–112.

[4] Gregoretti, I.V.; Lee, Y.M.; Goodson, H.V. Molecular evolution of the histone deacetylase family: functional implications of phylogenetic analysis. J. Mol. Biol. 2004, 338, 17–31.

[5] Preyat, N.; Leo, O. Sirtuin deacylases: a molecular link between metabolism and immunity. J. Leukoc. Biol. 2013, 93, 669–680.

[6] Matsushima, S.; Sadoshima, J. The role of sirtuins in cardiac disease. Am. J. Physiol. Heart Circ. Physiol. 2015 309(9), H1375–H1389.

[7] Mattagajasingh, I.; Kim, C.S.; Naqvi, A.; Yamamori, T.; Hoffman, T.A.; Jung, S.B.; DeRicco, J.; Kasuno, K.; Irani, K. SIRT1 promotes endothelium-dependent vascular relaxation by activating endothelial nitric oxide synthase. Proc. Natl. Acad. Sci. 2007, 104, 14855–14860.

[8] Potente, M.; Ghaeni, L.; Baldessari, D.; Mostoslavsky, R.; Rossig, L.; Dequiedt, F.; Haendeler, J.; Mione, M.; Dejana, E. SIRT1 controls endothelial angiogenic functions during vascular growth. Genes Dev. 2007, 21, 2644–2658.

[9] Li, X.; Zhang, S.; Blander, G.; Tse, J.G.; Krieger, M.; Guarente, L. SIRT1 deacetylates and positively regulates the nuclear receptor LXR. Mol. Cell 2007, 28, 91–106.

[10] Lagouge, M.; Argmann, C.; Gerhart-Hines Z.; Meziane, H.; Lerin, C.; Daussin, F.; Messadeq, N.; Milne, J. Lambert, P.; Elliott, P.; Geny, B.; Laakso, M.; Puigserver, P.; Auwerx, J. Resveratrol improves mitochondrial function and protects against metabolic disease by activating SIRT1 and PGC-1alpha, Cell 2006, 127, 1109–1122.

[11] Milne, J.C.; Lambert, P.D.; Schenk, S.; Carney, D.P.; Smith, J.J.; Gagne, D.J.; Jin, L.; Boss, O.; Perni, R.B.; Vu, C.B.; Bemis, J. E.; Xie, R.; Disch, J.S.; Ng, P.Y.; Nunes, J.J.; Lynch, A.V.; Yang, H.; Galonek, H.; Israelian, K.; Choy, W.; Iffland, A.; Lavu, S.; Medvedik, O.; Sinclair, D.A.; Olefsky, J.M.; Jirousek, M.R.; Elliott, P.J.; Westphal, C.H. Small molecule activators of SIRT1 as therapeutics for the treatment of type 2 diabetes, Nature 2007, 450, 712–716.

[12] Yamamoto, H.; Schoonjans, K.; Auwerx, J. Sirtuin functions in health and disease, Mol. Endocrinol. 2007, 21, 1745–1755.

[13] Zhao, X.; Allison, D.; Condon, B.; Zhang, F.; Gheyi, T.; Zhang, A.; Ashok, M.; MacEwan, R.I.; Qian, Y.; Jamison, J.A.; Luz, J.G. The 2.5 Å crystal structure of the SIRT1 catalytic domain bound to nicotinamide adenine dinucleotide (NAD+) and an indole (EX527 analogue) reveals a novel mechanism of histone deacetylase inhibition, J. Med. Chem. 2013,14, 963–969.

[14] Davenport, A.M.; Huber, F.M.; Hoelz, A. Structural and functional analysis of human SIRT1, J. Mol. Biol. 2014, 426, 526–541.

[15] Autiero, I.; Costantini, S.; Colonna, G. Human sirt-1: molecular modeling and structure-function relationships of an unordered protein, PLoS One 2009, 4, e7350.

[16] Sharma, A.; Gautam, V.; Costantini, S.; Paladino, A.; Colonna, G. Interactomic and pharmacological insights on human Sirt-1, Front. Pharmacol. 2012, 3, 40.

[17] Baur, J. A.; Pearson, K.J.; Price, N.L.; Jamieson, H.A.; Lerin, C.; Kalra, A. Prabhu, V.V.; Allard, J. S.; Lopez-Lluch, G.; Lewis, K.; Pistell, P.J.; Poosala, S.; Becker, K.G.; Boss, O.; Gwinn, D.; Wang, M.; Ramaswamy, S.; Fishbein, K.W.; Spencer, R.G.; Lakatta, E.G.; Le Couteur, D.; Shaw, R.J.; Navas, P.; Puigserver, P.; Ingram, D.K.; de Cabo, R.; Sinclair, D.A. Resveratrol improves health and survival of mice on a high-calorie diet. Nature 2006, 444, 337–342.

[18] Vu, C.B.; Bemis, J.E.; Disch, J.S.; Ng, P.Y.; Nunes, J.J.; Milne, J.C.; Carney, D.P.; Lynch, A.V.; Smith, J.J.; Lavu, S.; Lambert, P.D.; Gagne, D.J.; Jirousek, M.R.; Schenk, S.; Olefsky, J.M.; Perni, R.B. Discovery of imidazo[1,2-b]thiazole derivatives as novel SIRT1 activators. J. Med. Chem. 2009, 52, 1275–1283.

[19] Bemis, J. E.; Vu, C.B.; Xie, R.; Nunes, J. J.; Ng, P.Y.; Disch, J.S.; Milne, J.C.; Carney, D.P.; Lynch, A.V.; Jin, L.; Smith, J.J.; Lavu, S.; Iffland, A.; Jirousek, M.R.; Perni, R.B. Discovery of oxazolo[4,5-b] pyridines and related heterocyclic analogs as novel SIRT1 activators. Bioorg. Med. Chem. Lett. 2009, 19, 2350–2353.

[20] Mai, A.; Valente, S.; Meade, S.; Carafa, V.; Tardugno, M.; Nebbioso, A.; Galmozzi, A.; Mitro, N.; Fabiani, E.D.;. Altucci, L.; Kazantsev, A. Study of 1,4-dihydropyridine structural scaffold: discovery of novel sirtuin activators and inhibitors. J. Med. Chem. 2009, 52, 5496–5504.

[21] Howitz, K.T.; Zipkin, R.E. Compositions and methods for selectively activating human sirtuins. 2006, US20060014705A1.

[22] Milburn, M.; Milne, J.; Auwerx, J.; Argmann, C.; Lagouge, M.; Dipp, M. Methods and related compositions for treating or preventing obesity, insulin resistance disorders, and mitochondrial-associated disorders, 2007, US20070149466A1.

[23] Nayagam, V.M.; Wang, X.; Tan, Y.C.; Poulsen, A.; Goh, K.C.; Ng, T.; Wang, H.; Song, H.Y.; Ni, B.; Entzeroth, M. SIRT1 modulating compounds from high-throughput screening as anti-inflammatory and insulin-sensitizing agents. J. Biomol. Screen. 2006, 11, 959–967.

[24] Dai, H.; Case, A.W.; Riera, T.V.; Considine, T.; Lee, J.E.; Hamuro, Y.; Zhao, H.; Jiang, Y.; Sweitzer, S.M.; Pietrak, B.; Schwartz, B.; Blum, C.A.; Disch, J.S.; Caldwell, R.; Szczepankiewicz, B.; Oalmann, C.; Yee, Ng P.; White, B.H.; Casaubon, R.; Narayan, R.; Koppetsch, K.; Bourbonais, F.; Wu, B.; Wang, J.; Qian, D.; Jiang, F.; Mao, C.; Wang, M.; Hu, E.; Wu, J.C.; Perni, R.B.; Vlasuk, G.P.; Ellis, J.L. Crystallographic structure of a small molecule SIRT1 activator-enzyme complex. Nat. commun., 2015. 6, 7645.

[25] Sakkiah, S.; Arooj, M.; Lee, K.W.; Torres, J.Z. Theoretical approaches to identify the potent scaffold for human sirtuin1 activator: Bayesian modeling and density functional theory. Med. Chem. Res., 2014, 23, 3998–4010.

[26] Vyas, V.K.; Goel, A.; Ghate, M.; Patel, P., Ligand and structure-based approaches for the identification of SIRT1 activators, Chem. Biol. Interact. 2015, 228, 9–17.

[27] Gupta, A.K.; Bhunia, S.S., Balaramnavar, V.M.; Saxena, A.K. Pharmacophore modelling, molecular docking and virtual screening for EGFR (HER 1) tyrosine kinase inhibitors, SAR QSAR Env. Res. 2011, 22, 239–263.

[28] Gupta, A.K.; Saxena, S.; Saxena, M. Integrated ligand and structure-based studies of flavonoids as fatty acid biosynthesis inhibitors of Plasmodium falciparum, Bioorg. Med. Chem. Lett. 2010, 20, 4779–4781.

[29] Gupta, A.K.; Varshney, K.; Saxena, A.K. Toward the Identification of a Reliable 3D QSAR pharmacophore Model for the CCK2 Receptor Antagonism. J. Chem. Inf. Model. 2012, 52, 1376–1390.

[30] Khare, P.; Gupta, A.K.; Gajula, P.K.; Sunkari, K.Y., Jaiswal, A.K.; Das, S.; Bajpai, P., Chakraborty TK, Dube A, Saxena AK. Identification of novel s-adenosyl-l-homocysteine hydrolase inhibitors through homology-model-based virtual screening, synthesis, and biological evaluation, J. Chem. Inf. Model. 2012, 52, 777–779.

[31] Gupta, A.K.; Saxena, A.K. 3D-QSAR CoMFA and CoMSIA studies on a set of diverse α-adrenergic receptor antagonists. Med. Chem. Res. 2011, 20, 1455–1464.

[32] Gupta, A.K.; Saxena, A.K. Triple-layered QSAR studies on substituted 1,2,4-trioxanes as potential antimalarial agents: superiority of the quantitative pharmacophore-based alignment over common substructure-based alignment, SAR QSAR Env. Res. 2013, 24, 119–134.

[33] SYBYL 6.9, Tripos Inc., South Hanley Road, St. Louis, MO 63144, 1699.

[34] Gohda, K.; Mori, I.; Ohta, D.; Kikuchi, T.A.; CoMFA analysis with conformational propensity: an attempt to analyze the SAR of a set of molecules with different conformational flexibility using a 3D-QSAR method. J. Comput. Aided Mol. Des. 2000, 14, 265–275.

[35] Yap, C.W.; PaDEL-Descriptor: An open source software to calculate molecular descriptors and fingerprints. J. Comp. Chem. 2011, 32, 1466–1474.

[36] Wildman, S.A.; Crippen, G.M. Prediction of Physicochemical Parameters by Atomic Contributions. J. Chem. Inf. Comput. Sci. 1999, 39, 868–873.

[37] Todeschini, R.; Consonni, V. Molecular descriptors for chemoinformatics. Weinheim: Wiley VCH, 2009, 714–726.

[38] Wiener and Harry, Structural Determination of Paraffin Boiling Points, J. Am. Chem. Soc., 69(1947), pp. 17–20.

